# The Primary Cilium and its Hedgehog Signaling in Nociceptors Contribute to Inflammatory and Neuropathic Pain

**DOI:** 10.1101/2023.12.27.573420

**Authors:** Lindsey A. Fitzsimons, Larissa Staurengo-Ferrari, Oliver Bogen, Dioneia Araldi, Ivan J. M. Bonet, Ethan E. Jordan, Jon D. Levine, Kerry L. Tucker

## Abstract

The primary cilium, a 1-3 μm long hair-like structure protruding from the surface of almost all cells in the vertebrate body, is critical for neuronal development and also functions in the adult. As the migratory neural crest settles into dorsal root ganglia (DRG) sensory neurons elaborate a single primary cilium at their soma that is maintained into adult stages. While it is not known if primary cilia are expressed in nociceptors, or their potential function in the mature DRG neuron, recent studies have shown a role for Hedgehog, whose signaling demonstrates a dependence on primary cilia, in nociceptor sensitization. Here we report the expression of primary cilia in rat and mouse nociceptors, where they modulate mechanical nociceptive threshold, and contribute to inflammatory and neuropathic pain. When siRNA targeting *Ift88*, a primary cilium-specific intra-flagellar transport (IFT) protein required for ciliary integrity, was administered by intrathecal injection, in the rat, it resulted in loss of *Ift88* mRNA in DRG, and primary cilia in neuronal cell bodies, which was associated with an increase in mechanical nociceptive threshold, and abrogation of hyperalgesia induced by the pronociceptive inflammatory mediator, prostaglandin E_2_, and painful peripheral neuropathy induced by a neurotoxic chemotherapy drug, paclitaxel. To provide further support for the role of the primary cilium in nociceptor function we also administered siRNA for another IFT protein, *Ift*52. *Ift*52 siRNA results in loss of *Ift*52 in DRG and abrogates paclitaxel-induced painful peripheral neuropathy. Attenuation of Hedgehog-induced hyperalgesia by *Ift88* knockdown supports a role for the primary cilium in the hyperalgesia induced by Hedgehog, and attenuation of paclitaxel chemotherapy-induced neuropathy (CIPN) by cyclopamine, which attenuates Hedgehog signaling, suggests a role of Hedgehog in CIPN. Our findings support a role of nociceptor primary cilia in the control of mechanical nociceptive threshold and in inflammatory and neuropathic pain, the latter, at least in part, Hedgehog dependent.

## Introduction

The primary cilium, a single 1-3 μm long plasma membrane protrusion present on diverse cell types, contains a microtubule-based axoneme extending from a centriole-based basal body situated in the apical portion of the cytoplasm ^1,2^. A wide variety of clinical syndromes, termed “ciliopathies”, occur in patients with gene defects for proteins that localize to the cilium and its basal body ^3,4^. While changes in sensory neurons have been reported in patients with Bardet-Biedl syndrome ciliopathy ^5^, and pain has been reported by these patients ^5^, as well as by patients with Caroli disease ciliopathy ^6,7^, and in the majority of patients with the most common ciliopathy, autosomal dominant polycystic kidney disease ^8,9^, it is not known if primary cilia are present in nociceptors and their potential function related to pain.

During embryonic development, shortly after elaborating axons, dorsal root ganglion (DRG) neurons display a primary cilium at their soma ^10^. However, whether they retain this cilium in postnatal stages is controversial ^5,10^, and whether they occur in nociceptive sensory neurons is unknown. Given that neurons in the adult central nervous system (e.g. pyramidal neurons of the hippocampus) have been shown to depend upon primary cilia to modulate electrophysiological properties and gene transcription ^11^, the following hypothesis can be formulated: primary cilia are expressed by nociceptors, in which they modulate activation threshold and contribute to inflammatory and neuropathic pain, in the adult.

Binding of Hedgehog (Hh) ligands to the Patch1 receptor allows for translocation of the G protein-coupled receptor Smoothened (Smo) into the primary cilium, where it activates the Hh signal transduction pathway ^12,13^. Importantly, primary cilia are essential for signal transduction mechanisms of the canonical Hh pathway ^14^, which has been implicated in nociceptor sensitization ^15–20^. Thus, recent studies have reported a role for Hh signaling in the transduction and processing of nociceptive signals in both *Drosophila* ^21^ and rodents ^15–18,22,23^, morphine-induced hyperalgesia and tolerance ^15,18^, bone ^17^ and pancreatic ^16^ cancer pain, and chronic post-thoracotomy pain ^23^ (reviewed in ^20^). Therefore, in the present experiments we have explored the role of the primary cilium and cilium-dependent Hh signaling in baseline nociceptor function (i.e., setting mechanical nociceptive threshold) and in inflammatory and neuropathic pain.

## Materials and Methods

### Animals

All experiments using rats were performed on 300 to 400 g male Sprague-Dawley rats (Charles River Laboratories, Hollister, CA). Rats were housed 3 per cage under a 12-h light/dark cycle in a temperature- and humidity-controlled room at the University of California, San Francisco animal care facility. Food and water were available in home cages, *ad libitum*. Protocols for rat experiments were approved by the University of California, San Francisco, Institutional Animal Care and Use Committee, and adhered to the National Institutes of Health (NIH) Guidelines for the care and use of laboratory animals.

All studies on mice were performed with C57BL/6J male and female mice (The Jackson Laboratory, Bar Harbor, ME). Experiments in mice were conducted under a protocol approved by the Institutional Animal Care and Use Committee at the University of New England, in accordance with the Guide for the Care and Use of Laboratory Animals as adopted and promulgated by the NIH. Mice were housed 3-5 per cage under a 12-h light/dark cycle in a temperature- and humidity-controlled room in the animal care facility at the University of New England, Biddeford campus. Food and water were available in home cages, *ad libitum*.

### Immunohistofluorescence analysis of primary cilia in lumbar dorsal root ganglia (DRGs)

Rats were anesthetized with 5% isoflurane followed by perfusion through the left ventricle with 100 ml of cold phosphate buffered saline (PBS) containing 10U heparin/mL PBS, followed by 300 ml of methanol-buffered 10% formalin (Thermo Fisher, Waltham, MA). Bilateral L4 and L5 DRGs were dissected out, transferred into PBS containing 30% sucrose and 0.02% NaN_3_. These DRGs were then mounted in red tissue freezing medium (Electron Microscopy Sciences, Hatfield, PA) and 15 micrometers (μm) sections cut with a cryostat (Leica CM1900). Sections were then mounted on glass slides and used for immunohistofluorescence analyses.

Presence and length of primary cilia on DRG neurons were investigated using antibody staining of histological sections of adult rat DRGs and acutely-dissociated cultures prepared from surgically-isolated fresh mouse DRGs. Slides containing either rat DRG sections or coverslips with mouse DRG neuronal cultures were rehydrated and washed three times with 1X PBS, followed by a 30-min incubation in blocking buffer (1X PBS, 0.3% TritonX-100, 5% normal goat serum). Blocking buffer was then aspirated and primary antibodies, in antibody incubation buffer (1x PBS, 0.3% TritonX-100, 1% bovine serum albumin), were applied and allowed to incubate in a humid, dark chamber overnight at 4 °C. Slides were allowed to equilibrate to room temperature, and primary antibody solution aspirated. Slides were then washed three times with 1X PBS, secondary antibodies in antibody incubation buffer were applied and allowed to incubate at room temperature in a humid, dark chamber for 45 min. Slides were then washed five times with 1X PBS, followed by one rinse in a Coplin jar containing DI H_2_0, and finally cover slipped with DAPI mounting medium (Abcam, Waltham, MA) before imaging with a confocal microscope. The following primary antibodies were employed: rabbit anti-ARL13B (ADP-ribosylation factor-like 13B; ProteinTech, Rosemont, IL) 1:1000, mouse anti-ɣ-tubulin (clone GTU-88, Sigma-Aldrich, St. Louis, MO) 1:1000, goat anti-TrkA (tropomyosin receptor kinase A; R&D Systems, Minneapolis, MN) 1:300, rabbit anti-ACIII (adenylyl cyclase III; Encor, Gainesville, FL) 1:4000, mouse NeuN (RNA binding fox-1 homolog 3; ProteinTech) 1:1000, and Alexa 647-coupled IB4 (isolectin GS-IB_4_; Invitrogen) 1:500. The following secondary antibodies were employed: goat anti-rabbit IgG (H+L) (Alexa 488-coupled, Invitrogen, Waltham, MA) 1:1000, and goat anti-mouse IgG (H+L) (Alexa 555-coupled, Invitrogen) 1:1000.

### Imaging and data processing

Microscopy was conducted with a Leica Stellaris laser scanning confocal microscope utilizing Leica LAS X software version 4.6 and Plan Apo 40x/1.3NA and 63x/1.4NA objectives. 405, 488, 561, and 638 nm lasers with appropriate beam splitters (substrate or TD 488/561/633) were activated and emission ranges were selected for each fluorophore. Sequential scanning between frames was then activated at 1024×1024 resolution using frame average = 2. The channel containing the ciliary marker was then used to establish the appropriate z-volume. Optimal step size was established using the Nyquist Sampling Theorem calculated by LAS X. A z-stack of interest was then collected, titled, and saved for processing. Data was transferred as .lif files from the confocal software to FIJI. Channels were split individually, and stacks were projected to maximum intensity. LUT was set to grayscale for each of the channels to adjust brightness and contrast. LUTs were then set for cell nuclei in blue, cilia in green, basal body in red, and grayscale or yellow for neuronal markers.

### DRG neuron cultures

Acute cultures of dissociated DRG were prepared from C57BL/6J mice aged 4 – 6 weeks (The Jackson Laboratory), as described by Molliver and colleagues ^24^. In anesthetized mice, after transcardial perfusion of ice-cold phosphate-buffered saline solution, and removal of overlying dorsal skin, spinal ligaments and paraspinal muscles, laminectomy was performed with spring-loaded micro-iridectomy scissors and lumbar DRG were harvested. After desheathing, DRGs were pooled and treated, first with papain (Worthington Biochemical, Lakewood, NJ) for 10 minutes (min) at 37 °C, followed by a digestion with collagenase, type 2, (Worthington Biochemical) and dispase II (Roche, Mannheim, Germany) for 10 min at 37° C. After trituration with a fire-polished Pasteur pipette, neurons were plated onto individual 12-mm diameter round glass cover slips (Warner Instruments, Holliston, MA) that had been coated with poly-D-lysine and laminin and cultured in F12 medium with 5% fetal calf serum and penicillin-streptomycin (Thermo Fisher). Neurons were cultured for 2-5 days before fixation with 4% paraformaldehyde in a cytoskeletal-stabilizing buffer ^25^ at room temperature for 20 min, followed by immunohistofluorescence analysis.

### RNA extraction and reverse transcription quantitative real-time polymerase chain reaction (RT-qPCR)

Total RNA from rat L4 and L5 DRG was extracted using Trizol (Invitrogen) and the PureLink^TM^RNA Mini Kit (Invitrogen) according to the manufacturer’s instruction. The RNA concentration in each sample was determined with a spectrophotometer (Shimadzu, Santa Clara, CA). RNA was transcribed into cDNA using the iScript^TM^ Advanced cDNA Synthesis Kit for RT-qPCR (Bio-Rad, Hercules, CA). Real time PCR (Polymerase Chain Reaction) was performed with the SsoAdvanced Universal SYBR Green Supermix (Bio-Rad) and specific primers for IFT88 (intra-flagellar transport protein 88) (Bio-Rad, assay ID: qRnoCID0004310) and IFT52 (Bio-Rad assay ID: qRnoCID0002245) on the CFX 96^TM^ real-time PCR detection system (Bio-Rad). The PCR program was run with the following conditions: 2 min at 95 °C, followed by 40 cycles of 15 s at 95 °C, and 30 s at 60 °C. Each sample was run in triplicate. Change in gene expression was determined by the 2^-ΔΔCT^ method and expressed as relative change with respect to control level. Glycerinaldehyde-3-phosphate dehydrogenase (GAPDH) was used as a reference gene (Bio-Rad, assay ID: qRnoCID0057018).

### Pharmacological reagents

Prostaglandin E_2_ (PGE_2_), cyclopamine hydrate, and paclitaxel (purified from *Taxus yannanensis*) were purchased from Sigma-Aldrich. Recombinant rat Sonic hedgehog (Shh) was purchased from Abcam. Stock solutions of PGE_2_ (1 μg/μl) in absolute ethanol was diluted to 20 ng/μl) with 0.9% NaCl immediately before injection. The ethanol concentration of the final PGE_2_ solution was ∼2% and the injection volume 5 μl. Paclitaxel was prepared in absolute ethanol and polyethoxylated castor oil (Cremophor EL; 1:1; Sigma-Aldrich) and further diluted in saline, to a final concentration of 1 mg/kg). Cyclopamine hydrate was diluted in 0.9% NaCl containing 2% DMSO and was administered by intradermal (i.d.) or intra-ganglionic (i.gl.) routes of administration, both 10 μg), respectively. Recombinant Shh was reconstituted in 0.9% NaCl at 0.1 mg/ml and further diluted in 0.9% NaCl to be administered i.gl. or i.d. at 200 ng/animal. All dose selections were based on previous studies that established effectiveness at their targets ^17,26–28^.

### Intrathecal administration of siRNA

Small interfering (si) RNA was used to silence the mRNA expression of *Ift88* (mIFT88, Thermo Fisher, Ambion *in vivo* pre-designed siRNA, siRNA ID number S157132) and *Ift52* (mIFT52, Thermo Fisher Ambion *in vivo* predesigned siRNA pn4404010, siRNA ID number. S187205) *in vivo*, by intrathecal (i.t.) administration. A scrambled sequence was used as the negative control (nc-siRNA, Thermo Fisher, Ambion *in vivo* pre-designed negative control siRNA). siRNAs were mixed with PEI (*In Vivo* Jet-PEI; Polyplus Transfection) according to the manufacturer’s instructions. The N:P ratio (number of nitrogen residues of *In Vivo* Jet-PEI per DNA phosphate) used was 8 (1 μg of siRNA was mixed with 0.16 μl of *In Vivo* Jet-PEI). siRNA targeting IFT88, IFT52 and nc-siRNA were administered i.t. for 3 consecutive days. To administer siRNAs, rats were briefly anaesthetized with 2.5% isoflurane and a 30-gauge hypodermic needle inserted into the subarachnoid space, on the midline, between the L4 and L5 vertebrae. The i.t. site of injection was confirmed by the elicitation of a tail flick, a reflex that is evoked by accessing the subarachnoid space and bolus i.t. injection ^29^. This procedure produces not only reversible inhibition of the expression of the relevant proteins in DRG neurons but also modulation of nociceptive behavior ^27^.

### Intra-ganglion administration of pharmacological agents

Injection of compounds targeting the Hh signaling pathway into the L5 DRG was performed with a gingival needle (30-gauge) prepared as previously described ^30,31^, attached to a 50 μl Hamilton syringe by a short length of PE-10 polyethylene tubing; modification of the injecting needle decreases the risk of tissue damage when its tip penetrates the DRG. Rats were lightly anesthetized by inhalation of isoflurane, and the fur over the lower back shaved. The site of skin penetration was 1.5 cm lateral to the vertebral column and ∼0.5 cm caudal from a virtual line passing between the rostral edges of the iliac crests. To facilitate the insertion of the injecting needle through the skin, an initial cutaneous puncture was made with a larger (16-gauge) hypodermic needle. In sequence, the injecting needle was inserted through the puncture wound in the skin and oriented toward the region of the intervertebral space between the fifth and sixth lumbar vertebrae, until it touched the lateral region of the vertebrae. To reach the space between the transverse processes of the fifth and sixth lumbar vertebrae, small incremental movements of the needle tip were performed, until the resistance provided by the bone was diminished. When it reached the correct position, the tip of the needle felt “locked in place” (through the intervertebral space) and a flinch of the ipsilateral hindpaw was observed ^30^, indicating that it had penetrated the DRG of the fifth lumbar spinal nerve ^31^. The procedure, starting from the beginning of anesthetic inhalation, until withdrawal of the injection needle, takes ∼3 min. Animals regained consciousness ∼2 min after anesthesia was discontinued. Importantly, no change in the nociceptive mechanical paw-withdrawal threshold of the ipsilateral hindpaw, after a single injection or daily repeated injections in the L5-DRG, was observed, and immunofluorescence analysis of the L5-DRG after injections has shown no signs of cell damage ^30^. Likewise, no significant change in the mechanical nociceptive threshold in the hindpaw, after i.gl. injection of vehicle (single or repeated injections), was observed, indicating a lack of injection-induced damage (*PMID: 23401543*). This technique has been used by us and others (PMIDs: 25878283, 23401543, 25489099, 29626652, 32523543, 23776243, 30537520).

### Intradermal administration of pharmacological agents

For the i.d. injection of cyclopamine and recombinant Shh, a hypotonic shock (2 μl of distilled water, separated in the syringe by an air bubble) preceded their injection, to facilitate entry of membrane-impermeable compounds into nerve terminals^32–34^ on the dorsum of the hind paw. In these experiments, a total volume of 5 μl was injected i.d. on the dorsum of the hind paw.

### Measuring mechanical nociceptive threshold

Mechanical nociceptive threshold was quantified using an Ugo Basile Analgesymeter (Stoelting, Wood Dale, IL), to perform the Randall–Selitto paw withdrawal test ^35–37^. This device applies a mechanical force that increases linearly with time, to the dorsum of the rat’s hind paw. Rats were placed into cylindrical acrylic restrainers with lateral ports that allow access to the hind paw, as previously described ^38^, to acclimatize them to the testing procedure. Mechanical nociceptive threshold is defined as the force in grams at which a rat withdraws its paw. Baseline threshold is defined as the mean of 3 readings taken before injection of test agents; each experiment was performed on different groups of rats. Data are presented as mechanical nociceptive threshold (in grams) and as percentage change from preintervention baseline.

### Paclitaxel chemotherapy-induced peripheral neuropathy (CIPN) in the rat

Using an established rat CIPN model, paclitaxel was injected intraperitoneally (i.p.) every other day for a total of 4 doses (1 mg/kg × 4 i.p.); control animals received the same volume of vehicle with proportional amounts of Cremophor EL and ethanol diluted in saline ^28,39^. Presence of paclitaxel-induced painful peripheral neuropathy was confirmed by demonstrating a decrease in mechanical nociception threshold in the Randall-Selitto paw withdrawal test ^28,40–42^.

### Inflammatory pain model: PGE_2_-induced mechanical hyperalgesia in the rat

PGE_2_ (100 ng) was injected i.d., in a volume of 5 µl, using a 30-gauge hypodermic needle attached to a microsyringe (Hamilton, Reno NV). PGE_2_-induced hyperalgesia was quantified as the percentage change in mechanical nociceptive threshold, calculated from the baseline, pre-injection, mechanical nociceptive threshold versus the threshold after injection of the test agent. The dosing protocol and timing of mechanical nociceptive threshold assessment were based on previous studies ^26,43^.

### Statistical analyses

In each behavioral experiment, only one hind paw per rat was used. In these experiments, data are presented as percentage change from baseline mechanical nociceptive threshold. Repeated-measures one-way analysis of variance (ANOVA), followed by the Bonferroni *post hoc* multiple comparison test or the Student t test was used for statistical analyses. Prism 8.0 (GraphPad Software, San Diego, CA) was used for the graphics and to perform statistical analyses; P < 0.05 was considered statistically significant. Data are presented as mean ± SEM.

## Results

### Adult nociceptors elaborate soma primary cilia, *in vivo* and *in vitro*

Primary cilia in adult DRG neurons were detected using immunohistofluorescence staining with confocal microscopy. Adult rat DRGs (**Fig. 1A-D, Suppl. Fig. 1B**), and cultures of neurons from acutely-dissociated adult mouse DRGs (**Suppl. Fig. 1A**), were labeled with an antibody recognizing the ciliary protein Arl13b (ADP-ribosylation factor-like 13B) ^44^, to identify the plasma membrane surrounding the central ciliary axoneme, and an antibody recognizing ɣ-tubulin to label the basal body from which the primary cilium grows ^45,46^. To identify the cells expressing these ciliary markers as neurons, we employed antibodies recognizing Fox-3/NeuN (**Fig. 1A)**, TrkA (**Fig. 1B**) or calcitonin gene-related peptide (CGRP) (**Fig. 1C**). We also identified neurons binding isolectin IB4, which labels non-peptidergic nociceptors (**Fig. 1D**). In DRG cultures neurons were identified by their double birefringent phase-contrast appearance and isolectin B4 (IB4) reactivity (**Suppl. Fig. 1A**). Clear indication of a primary cilium was found on many neurons in the rat DRG and elaborated by cultured mouse DRG neurons. In rat DRGs, the large number of satellite cells surrounding neurons are also ciliated (**Suppl. Fig. 1B**), with cilia that appear to be oriented facing neuronal cell bodies.

**Figure 1.**
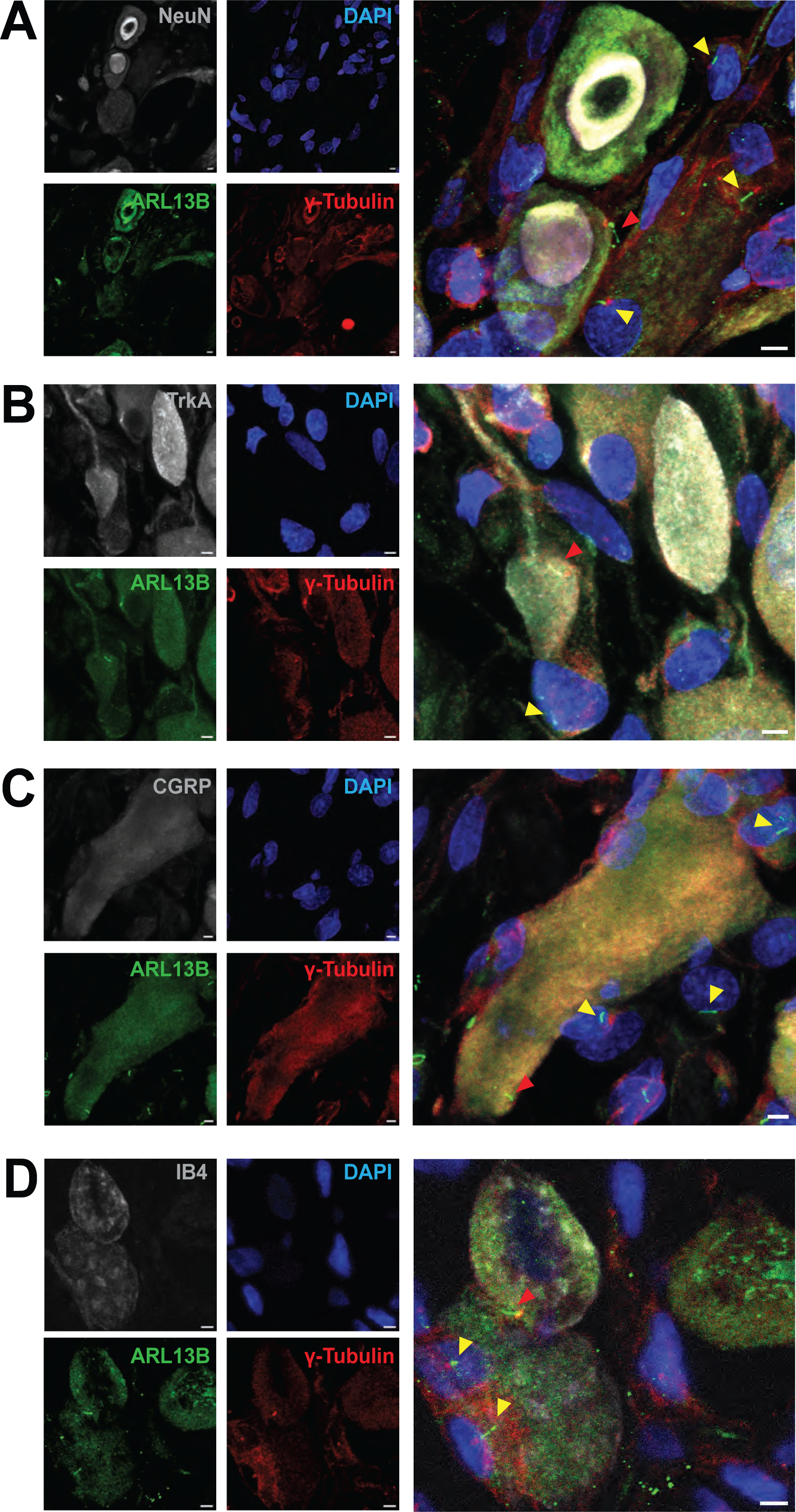
Rat DRG neurons elaborate primary cilia, *in vivo*. (**A-D**) Immunohistofluorescence analysis of adult rat DRGs *in vivo*, using confocal microscopy. Histological sections were labeled with antibodies recognizing ARL13B (**A-D,** green), ɣ-tubulin (**A-D,** red), and Fox-3 (**A,** NeuN, greyscale), TrkA (**B,** greyscale), or CGRP (**C,** greyscale). (**D**) IB4 staining is indicated in greyscale. (**A-D**) Cell nuclei are marked by DAPI (blue). Red and yellow arrowheads indicate neuronal and non-neuronal primary cilia, respectively. (**A-D**) Scale bar 3 µm.

### Targeted knockdown of *Ift88* by siRNA attenuates *Ift88* in rat DRG, leading to a loss of primary cilia and a decrease in the average length of retained cilia

Two classes of IFT protein complexes, B and A, are responsible for anterograde and retrograde trafficking of proteins into and out of the primary cilium, respectively ^47^. *Ift88* forms an important component of the B complex ^48^. Primary cilia in DRG neurons can be disrupted through the targeted inactivation of genes encoding proteins essential for maintenance of primary cilia structure, such as *Ift88*, an approach that we have employed successfully ^49–52^. In the present experiments male rats were treated i.t. with siRNA targeting *Ift88,* in a dose of 2, 5 or 10 μg/day, for 3 consecutive days (Fig. 3A-D). Because rats treated with the highest dose of siRNA showed the strongest behavioral phenotype, L4 and L5 DRGs were harvested 7 days after the last injection of *Ift*88 siRNA, from euthanized rats, to prepare mRNA, or rats were transcardially perfused with 4% paraformaldehyde to prepare tissue for immunohistofluorescence analysis. Total RNA, isolated from L4 and L5 lumbar DRGs, from rats treated with 10 μg siRNA, was used to generate cDNA through reverse transcription. Quantitative RT-PCR, performed on the resultant cDNA revealed a significant decrease in *Ift88* expression in animals treated with the 10 μg dose of siRNA targeting *Ift88* (**Fig. 2A**). To visualize and quantify the primary cilium at the single-cell level, DRGs were harvested from paraformaldehyde-perfused rats, 7 days after the final injection of the 10 μg dose of siRNA targeting *Ift88*. Then 15-μm thick sections of cryo-embedded L4 and L5 DRGs were labeled with antibodies recognizing Arl13b and ɣ-tubulin, followed by confocal microscopy (**Fig. 2B**). Counting of cells with and without cilia, and length measurement of remaining cilia revealed a significant decrease in both the prevalence of cilia in DRG cells (**Fig. 2C**) as well as the average length of remaining cilia (**Fig. 2D**), in animals treated with siRNA targeting *Ift88*, compared to controls treated with a negative siRNA.

**Figure 2.**
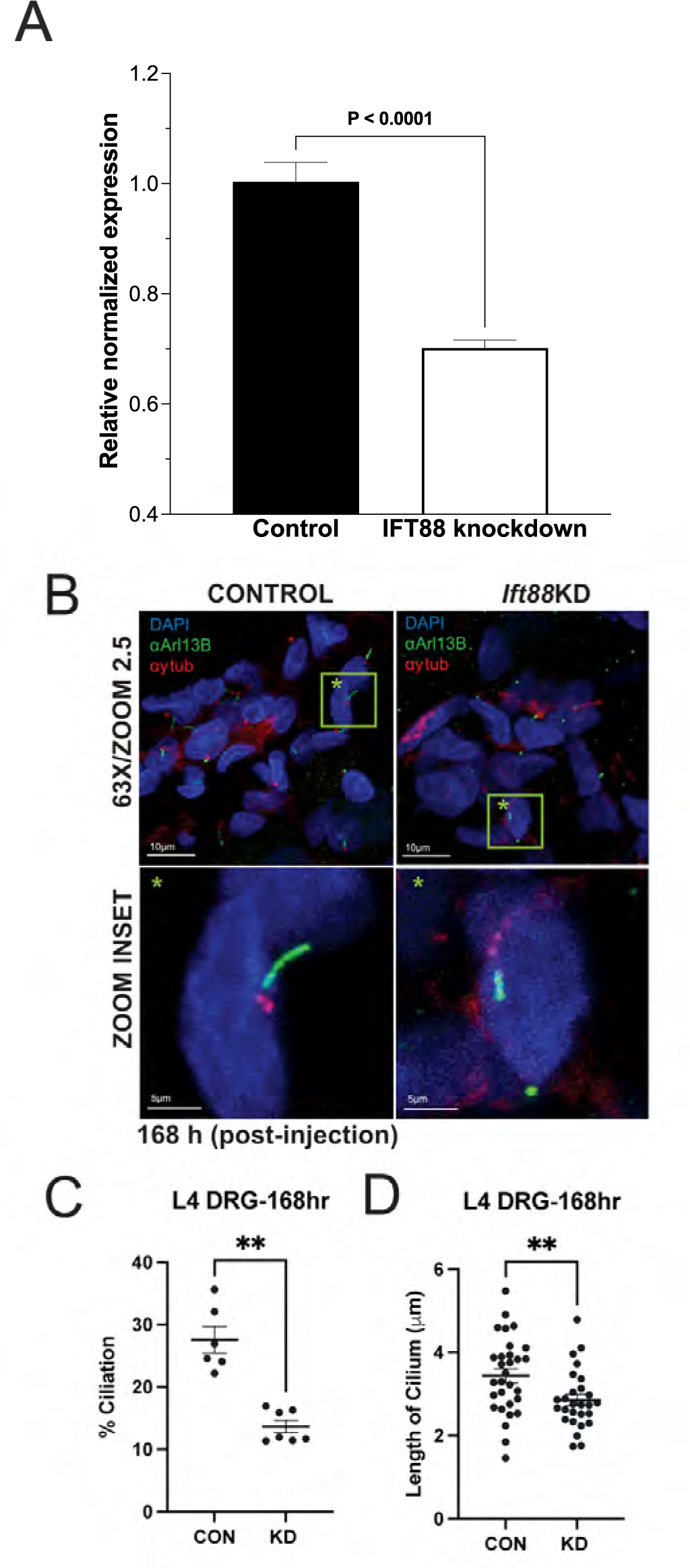
Targeted knockdown of Ift88 in rat DRG leads to a loss of primary cilia and a decrease in the average length of retained cilia. **(A)** siRNA-mediated knockdown of *Ift88*. The histogram depicts the level of *Ift88* expression in rats treated with either a control siRNA (control) or the siRNA targeting *Ift88* (Ift88 knockdown). Gene expression was determined with SYBR green RT-PCR and values (mean ± 95% CI) represent expression relative to GAPDH (the housekeeping gene). Rats treated with siRNA targeting *Ift88* demonstrate a significant reduction in *Ift88* mRNA in lumbar DRG, compared to those treated with the control siRNA (unpaired two-tailed Student’s t-test, P <0.0001, t(10)=7.88, n=6). One way ANOVA with Tukey comparison was used to analyze for differences between groups. (**B**) Confocal immunohistofluorescence images of L4 rat DRGs stained for the ciliary axoneme (αArl13b, green), the basal body (anti-ɣ-tubulin, red), and the nucleus (DAPI, blue). Left panels, control siRNA. Right panels, *Ift88*-targeted siRNA. Lower panels represent magnifications of the green boxes (green star) indicated in the upper panels. Magnification/zoom factor indicated. Scale bar 100 and 10 µm (top and bottom panels, respectively). (**C, D**) Quantification of percentage ciliation (**C**) and ciliary length (**D**) in micrometers (μm) in L4 rat DRGs (rDRG) injected with control (CTL) vs *Ift88*-targeted (KD) siRNA. **: P < 0.01.

**Figure 3.**
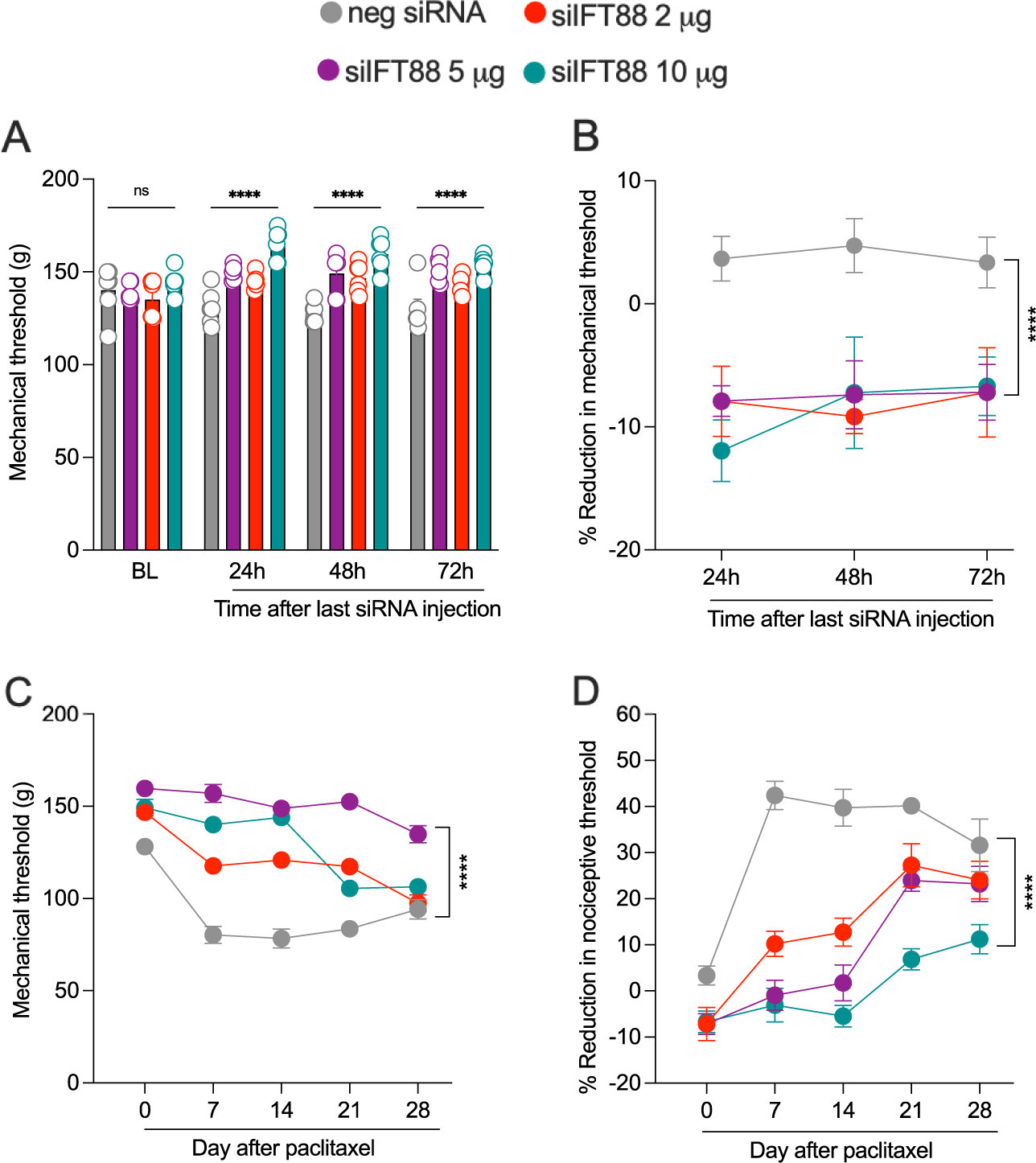
Effect of siRNA targeting *Ift88* on mechanical nociceptive threshold and paclitaxel-induced hyperalgesia (CIPN). Male rats were treated i.t. with one of 3 doses of siRNA (2, 5 or 10 µg/20 µl) targeting *Ift88* or negative control siRNA, 10 mg/20ml for 3 consecutive days. Mechanical nociceptive threshold was evaluated before siRNA injection was started and again at 24, 48, and 72 h after treatment was finished. 72 h after the last siRNA injection (day 0), paclitaxel (1 mg/kg, i.p.) was administered (days 0, 2, 4 and 6) and the mechanical nociceptive threshold evaluated on days 0, 7, 14, 21, and 28. (**A,B**) Administration of siRNA targeting *Ift88* increases mechanical nociceptive threshold when compared with the vehicle control group. (**A**) Mechanical nociceptive threshold is presented as absolute values in grams. Data is reported as mean ± SEM, treatment *F _(3,80)_* = 34.54, ****P <0.0001: neg siRNA vs si*Ift88* 2 µg, neg siRNA vs si*Ift88* 5 µg or neg siRNA vs si*Ift88* 10 µg. (**B**) Mechanical nociceptive threshold is represented as percentage change from baseline threshold. Data shown as mean ± SEM, treatment *F _(3,60)_* = 15.80, ****P <0.0001: neg siRNA vs si*Ift88* 2 µg, neg siRNA vs si*Ift88* 5 µg or neg siRNA vs si*Ift88* 10 µg. (**C,D**) The magnitude of paclitaxel-induced hyperalgesia was markedly attenuated in rats treated with siRNA targeting m*Ift88*. (**C**) Mechanical nociceptive threshold is presented as absolute values in grams. Data shown as mean ± SEM, treatment *F _(3,100)_* = 192.0, ****P <0.0001: neg siRNA vs si*Ift88* 2 µg, neg siRNA vs si*Ift88* 5 µg or neg siRNA vs si*Ift88* 10 µg. (**D**) Data shown as mean ± SEM, treatment *F _(3,100)_* = 77.50, ****P <0.0001: neg siRNA vs si*Ift88* 2 µg, neg siRNA vs si*Ift88* 5 µg or neg siRNA vs si*Ift88* 10 µg. Two-way repeated-measures ANOVAs were used to compare negative control siRNA and siRNA targeting *Ift88* mRNA groups, n = 6 paw in each group.

### Effect of *Ift88* siRNA on baseline mechanical nociceptive threshold and pain associated with paclitaxel CIPN

Male rats were treated i.t. with siRNA targeting *Ift88*, 2, 5 or 10 μg/day for 3 consecutive days. Twenty-four, 48 and 72 h after the last administration of siRNA, nociceptive threshold was evaluated by the Randall-Selitto paw-withdrawal test. At 24, 48 and 72 h, nociceptive threshold was significantly increased compared to rats receiving the same dose of a negative control siRNA (**Fig. 3A and B)**.

We next employed paclitaxel CIPN as a preclinical model of neuropathic pain, to evaluate the role of the nociceptor primary cilium in neuropathic pain. A group of male rats were treated with siRNA targeting *Ift88* (2, 5 or 10 μg/day, i.t.) for 3 consecutive days. Seventy-two hours after the last administration of siRNA, paclitaxel (1 mg/kg, i.p.) was administered on days 0, 2, 4 and 6. Mechanical nociceptive threshold was evaluated before siRNA treatment was started (baseline), and again on days -3, -2, -1, 3, 5, 7,14, 21 and 28 dose after the last dose of paclitaxel, from day 3 of behavioral testing, with the highest dose of *Ift88* siRNA bringing animals essentially back to baseline values, as compared with rats receiving an i.t. dose of a negative control siRNA (**Fig. 3C and D)**.

### Effect of *Ift52* siRNA on CIPN pain

To independently validate the role of primary cilia in nociceptor function we evaluated the role of a second intra-flagellar protein, *Ift52,* in paclitaxel CIPN. Male rats were treated with siRNA targeting *Ift52* (10 μg, i.t., for 3 consecutive days). Seventy-two hours after the last administration of *Ift52* siRNA rats were euthanized and L4 and L5 DRG harvested, for preparation of RNA. Total RNA isolated from L4 and L5 DRGs was used to generate cDNA through reverse transcription. Quantitative RT-PCR performed upon the resultant cDNA revealed a significant decrease in *Ift52* expression in animals treated with siRNA targeting *Ift52* (**Fig. 4A**). We again employed paclitaxel CIPN as our neuropathic pain model, to evaluate the role of the nociceptor primary cilium in neuropathic pain. Male rats were treated with siRNA targeting *Ift52* for 3 consecutive days in a dose of 10 μg/day (i.t.). Seventy-two hours after the last siRNA injection, paclitaxel (1 mg/kg, i.p.) was administered on days 0, 2, 4 and 6. Mechanical nociceptive threshold was evaluated before siRNA treatment was started (baseline), and again on days -3, -2, -1, 3, 5, 7,14, 21 and 28 days after the last dose of paclitaxel. Similar to the effect of *Ift88* siRNA, the siRNA targeting *Ift52* reduced paclitaxel-induced hyperalgesia from day 3 of behavior testing, as compared with rats receiving an i.t. dose of a negative control siRNA **(Fig. 4B and C)**.

**Figure 4.**
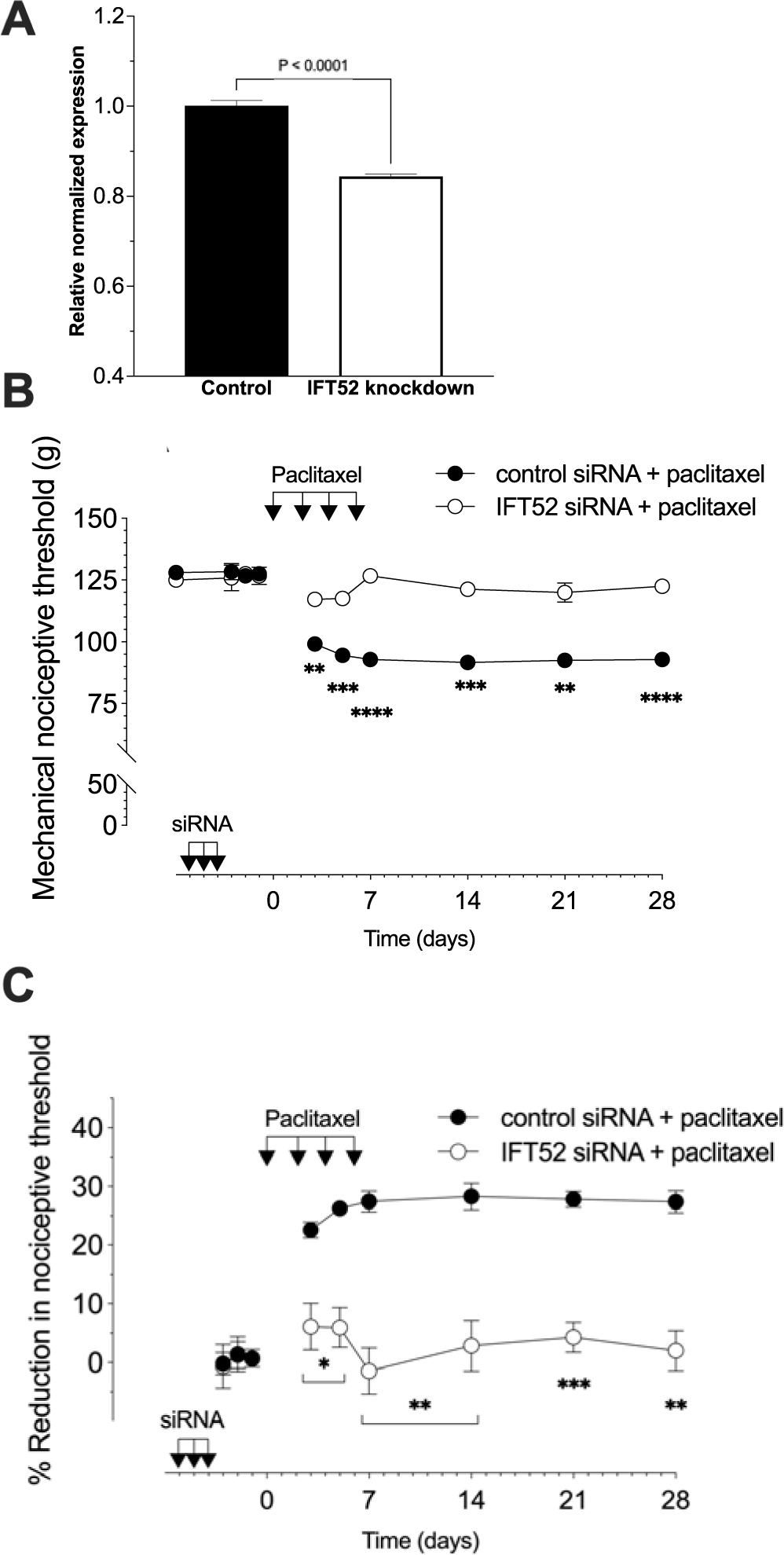
Role of IFT52 in paclitaxel-induced mechanical hyperalgesia. **(A)** siRNA-mediated knockdown of *Ift52* mRNA. The histogram depicts the level of *Ift52* mRNA in rats treated with either control siRNA or siRNA targeting *Ift52*. Gene expression was determined with SYBR green RT-PCR and values (mean ± 95% CI) represent expression relative to GAPDH (the housekeeping gene). Rats treated with siRNA targeting *Ift52* demonstrate a significant reduction in *Ift52* mRNA levels in their lumbar DRG compared to those treated with the control siRNA (unpaired two-tailed Student’s t-test, P <0.0001, t(10)=11.21, n=6). (**B,C**) Male rats were treated i.t. with siRNA targeting *Ift52* (10 µg/day) for 3 consecutive days. 72 h after the last siRNA injection, paclitaxel (1 mg/kg, i.p.) was administered on days 0, 2, 4 and 6. Mechanical nociceptive threshold was evaluated before siRNA treatment was started (baseline), and again on days -3, -2, -1, 3, 5, 7, 14, 21 and 28 days after paclitaxel injection. (**B**) Magnitude of hyperalgesia is expressed as absolute values of mechanical nociceptive threshold, in grams. *Ift52* siRNA attenuates paclitaxel induced CIPN. Data are expressed as means ± SEM, *n* = 6 paws in each group. F_(9,90)_ = 29.38, ****P <0.0001, ***P <0.0002, **P <0.0028; when *Ift52* siRNA group was compared with control siRNA group; two-way repeated-measures ANOVA followed by Bonferroni multiple comparison test. (**C**) The magnitude of hyperalgesia expressed as percentage reduction from the baseline mechanical nociceptive thresholds. *Ift52* siRNA attenuates paclitaxel induced CIPN. Data are expressed as means ± SEM, *n*=6 paws in each group. F_(8,80)_=12.87, ***P = 0.0005 **P <0.0089 *P = 0.0143; when the *Ift52* siRNA group was compared with the control siRNA group; two-way repeated-measures ANOVA followed by Bonferroni multiple comparison test.

### Primary cilium-dependence of the pronociceptive effect of Sonic hedgehog

While many effects of Hh are primary cilium-dependent^14,53^, the role of the primary cilia in Hh-induced pain has not been studied. To test for the dependence of the pronociceptive effects of Shh on the primary cilium, male rats were treated with siRNA targeting *Ift88*, in a dose of 10 μg/day for 3 consecutive days. Seventy-two hours after administration of the last dose of *Ift88* siRNA, rats received an i.gl. (**Fig. 5A**) or i.d. (**Fig. 5B**) injection of recombinant Shh (200 ng) and nociceptive threshold was evaluated. Treatment of rats with siRNA for *Ift88*, reduced hyperalgesia by both i.gl. and i.d. Shh (**Fig. 5A and B**).

**Figure 5.**
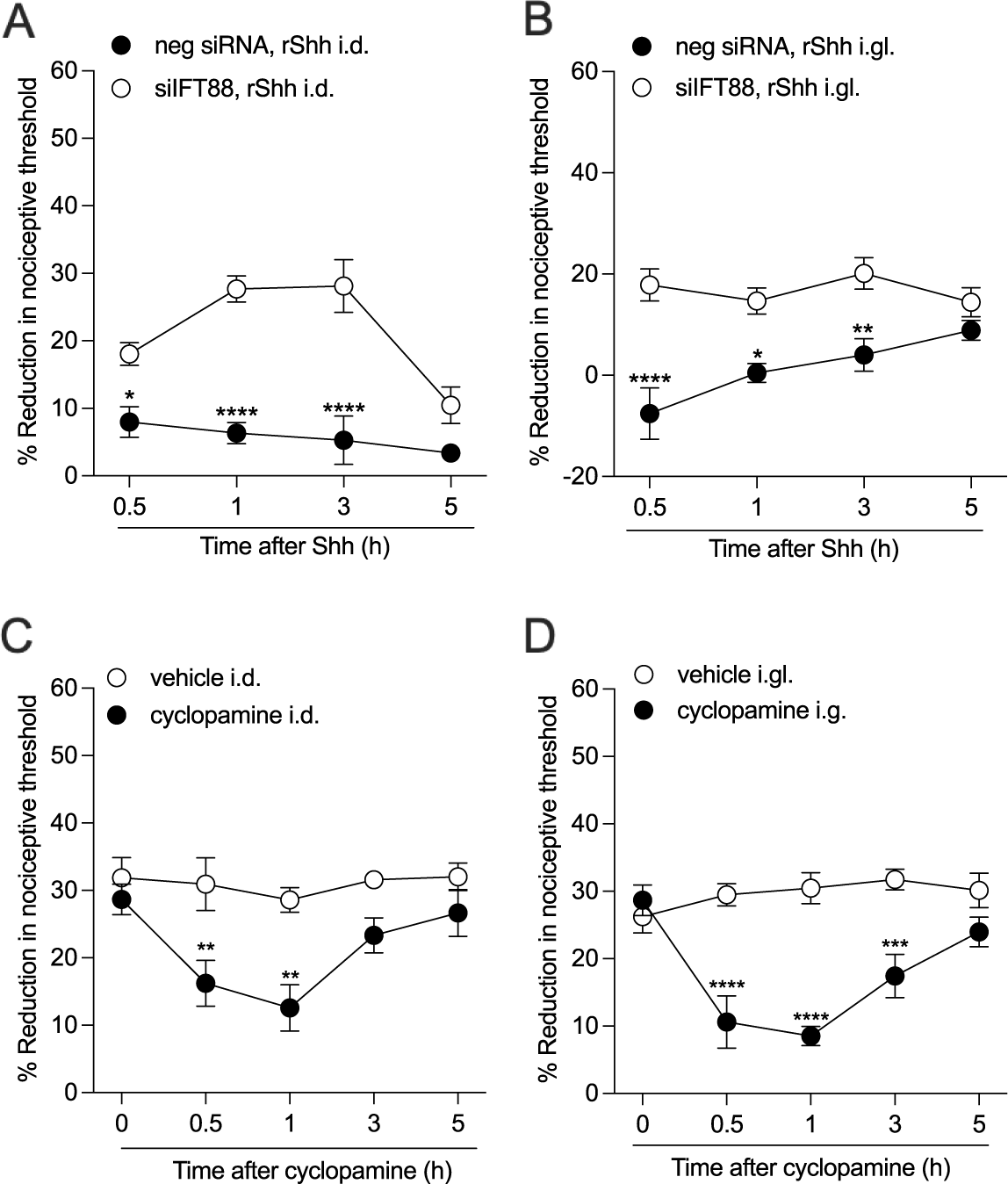
Pronociceptive effect of Shh is primarily cilium-dependent. (**A,B**) Male rats were treated i.t. with siRNA targeting *Ift88* (10 µg/day) for 3 consecutive days. 72 h after the last siRNA injection, rats received i.d. or i.gl. injections of recombinant Shh (200 ng) and mechanical nociceptive threshold was assessed at 0.5, 1, 3, and 5 h after injection. (**A**) The hyperalgesic effect of Shh injected i.d. was significantly reduced when the expression of Ift88 was downregulated through siRNA-mediated knockdown. Data shown as mean ± SEM, treatment *F _(1,40)_* = 73.26, time F _(4,50)_ = 6.861, ****P <0.01: siIFT88, Shh i.d. vs siRNA neg, Shh i.d.. (**B**) The hyperalgesic effect of i.gl. Shh was also significantly reduced when the expression of Ift88 was downregulated with siRNA. Data shown as mean ± SEM, treatment *F _(1,40)_* = 48.13, time F _(3,40)_ = 2.286, ****P <0.01: IFT88 siRNA, Shh i.gl. vs siRNA neg, Shh i.gl. Two-way repeated-measures ANOVA followed by Bonferoni’s test, n=6 paws in each group. (**C, D**) Male rats received paclitaxel every other day for a total of 4 injections, on days 0, 2, 4, and 6 (1 mg/kg × 4, i.p.). Seven days after the first administration of paclitaxel, rats were treated with i.d. (10 µg/5 µl) or i.gl. (10 µg/5 µl) cyclopamine. As a control, the contralateral paw or contralateral lumbar DRG received vehicle (saline plus DMSO 2%). Mechanical nociceptive threshold was evaluated before and 7 days after paclitaxel injection (before cyclopamine injection, ∼0h), and again 0, 0.5, 1, 3, and 5 h after i.d. or i.gl. cyclopamine injection. (**C**) Paclitaxel-induced hyperalgesia was markedly attenuated in the male rats treated i.d. with cyclopamine. Data shown as mean ± SEM, treatment *F _(1,50)_* = 28.28, time F _(4,50)_ = 4.154, **P <0.01: cyclopamine i.d. vs vehicle i.d. groups. (**D**) The magnitude of paclitaxel-induced hyperalgesia was markedly attenuated in male rats treated with i.gl. cyclopamine. Data shown as mean ± SEM, treatment *F _(1,50)_* = 57.38, time F _(4,50)_ = 4.748, **P <0.01: cyclopamine i.d. vs vehicle i.d..

To investigate the role of Hh signaling in nociceptor sensitization, we again employed the CIPN model of neuropathic pain induced by paclitaxel. To induce CIPN, male rats were treated with paclitaxel (1 mg/kg x 4, i.p.) on days 0, 2, 4 and 6. On day 7, at which time hyperalgesia was fully established, rats were treated with cyclopamine (10 μg) i.gl. (**Fig. 5D**) or i.d. (**Fig. 5C**), which inhibits Hh signaling by binding to the Smo co-receptor ^54^. Nociceptive threshold was evaluated 30 min and, 1 and 3 h after cyclopamine treatment (N=6). Administration of cyclopamine, by both the i.gl. and i.d. route, attenuated paclitaxel-induced hyperalgesia (**Fig. 5A and B**).

### siRNA targeting *Ift88* attenuates inflammatory pain

Finally, we examined the role of nociceptor primary cilia in inflammatory pain, as produced by i.d. injection of PGE_2_, a prototypical direct-acting pronociceptive inflammatory mediator^55–57^. Rats were treated i.t. with siRNA targeting *Ift88* (10 μg/day, i.t.) for 3 consecutive days. 72 h after the final siRNA injection, PGE_2_ was administered on the dorsum of the hindpaw (100ng/5μl/paw, i.d.), at the site of nociceptive threshold testing. Nociceptive threshold was evaluated before siRNA treatment was started (baseline), again 72 h after the last siRNA treatment, and 30 minutes after i.d. injection of PGE_2_ (**Fig. 6**). As was observed with baseline thresholds in the control rat (**Fig. 3A, B**), siRNA targeting of *Ift88* led to an increase in the baseline nociceptive threshold (**Fig. 6A**). Treatment with *Ift88* siRNA also resulted in a marked reduction in PGE_2_-induced mechanical hyperalgesia, evaluated after the last dose of *Ift88* siRNA (Fig. 6B), and 72 h after the last dose of *Ift88* siRNA (**Fig. 6C**).

**Figure 6.**
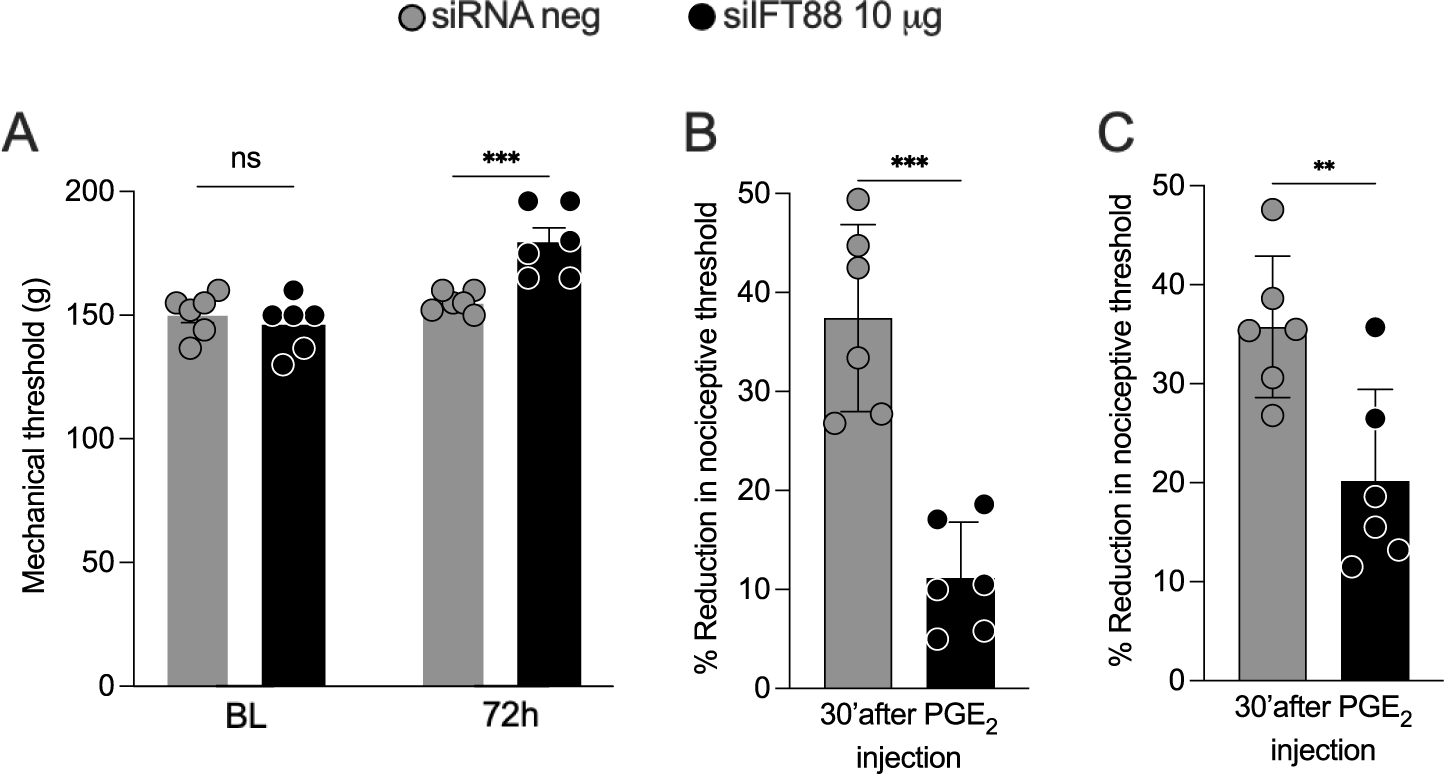
*Ift88* siRNA attenuates hyperalgesia induced by i.d. PGE_2_. Male rats were treated i.t. with siRNA for IFT88 mRNA for 3 consecutive days in a dose of 10 µg/day. 72 h after the last siRNA injection, PGE_2_ was administered i.d. (100 ng/5 µl/paw). Mechanical nociceptive threshold was evaluated before siRNA treatment was started (baseline), and again 72 h after the last siRNA treatment, and then 30 min after PGE_2_. (**A**) As shown in Figure 3, siRNA targeting Ift88 led to a significant increase in baseline nociceptive threshold, expressed in grams. Data shown as means ± SEM, Time F _(1,20)_ = 21.73 ***P < 0.0001: siIFT88 vs baseline or vs siRNA neg control. Two way-repeated measures ANOVA followed by Bonferroni’s test, n = 6 paws of each group. (**B, C**) Treatment with *Ift88* siRNA also resulted in a substantial reduction in hyperalgesia induced by i.d. PGE_2._ (**B**) The magnitude of hyperalgesia is expressed as percentage reduction from the baseline mechanical nociceptive threshold before siRNA treatment. Data shown as mean ± SEM, unpaired t-test: t = 5.846, df = 10; ***P < 0.001: siIFT88 vs siRNA neg control, n = 6 paws of each group. (**C**) The magnitude of hyperalgesia is expressed as % reduction from the baseline mechanical nociceptive thresholds read 72 h after the last treatment with siRNA. Data are expressed as mean ± SEM, unpaired t-test: t = 3.260, df=10; **P < 0.01: siIFT88 vs siRNA neg, n = 6 paws of each group.

## Discussion

In contrast to many cell types in the central nervous system, ciliation of DRG neurons has only recently been reported ^5,10,58^. The reason for this derives, in part, from the large size of the some in DRG neurons, which makes prohibitive the detection, through its serial sectioning for transmission electron microscopy, of a single 1-3 μm protrusion into the extracellular space from a spherical cell body of diameter as large as 60 μm. In addition, the close proximity of the large number of primary cilia expressing satellite cells ensheathing each neuronal cell body (**Supp. Fig. 1B)** make it difficult to ascribe neuronal ownership of a given primary cilium. While two groups have reported that neurons from acutely dissociated DRG, prepared from adult mice, bear a primary cilium ^5,58^, using similar preparations, another group has observed only few ciliated neurons ^10^. In the present experiments we have confirmed the presence of primary cilia on DRG neurons as well as shown, for the first time, based on the expression of characteristic proteins, that the ciliated DRG neurons include diverse populations of nociceptors. To our knowledge, this is the first demonstration of primary cilia on DRG neurons, *in situ,* within the milieu of the adult ganglion, as well as establishing that ciliated DRG neurons include heterogeneous populations of nociceptors, raising the possibility that primary cilia contribute to nociceptor function, including playing a role in inflammatory and neuropathic pain syndromes.

Primary cilia are often considered to act as antennae, an integral part of its cell’s sensory apparatus ^59–61^. Therefore, after establishing that nociceptors express primary cilia, we tested the hypothesis that they are involved in the transduction of noxious stimuli and contribute to nociceptor sensitization in inflammatory and neuropathic pain. We found that siRNA targeting *Ift88*, an integral ciliary protein necessary for the intra-flagellar transport of proteins essential for ciliary structure and function ^62,63^, attenuates the primary cilia in cells in the DRG, produces an elevation of mechanical nociceptive threshold and attenuates inflammatory and neuropathic pain. The mechanism by which the primary cilium, located at the nociceptor cell soma, contributes to the transduction of mechanical stimuli and nociceptor sensitization, in the setting of inflammatory and neuropathic pain at the peripheral terminal of the nociceptor, a long distance from the cell body, and whether the nociceptor primary cilium contributes to the detection of other noxious stimuli (e.g., noxious heat and cold, and pain-produced by noxious chemicals such as capsaicin) remains to be established. As the central terminal of the nociceptor, located in the dorsal horn of the spinal cord, is involved in neurotransmission, to second order neurons in pain circuitry, a possible role of primary cilia in pain neurotransmission, established in the central nervous system ^64–66^, should also be considered in nociceptors.

A precedent for cilium-dependent long-range control of biological events at remote terminal processes is afforded by examination of axonal pathfinding in embryonic neurons in the chick and the mouse. Axonal growth cones of neurons crossing the ventral midline of the spinal cord were found to employ retrograde transport of midline-derived Shh for activation of Hh signaling cascades at the primary cilium on the cell body of the same neuron, localized at the neuronal soma, hundreds of cell diameters away ^67^. Similarly, retinal ganglion cells have been reported to use anterograde transport of Shh to the optic chiasm, where secretion could direct axonal outgrowth ^68^. However, the timeframe of nociceptive modulation seen following peripheral application of Shh and cyclopamine in the present experiments would seemingly preclude the involvement of such retrograde signaling. Thus, alternative explanations appear to be required. Shh enhances calcium oscillations in embryonic neurons ^69^ and neural stem cells ^70^, and calcium signaling can propagate rapidly along axons ^71,72^, while primary cilia are specialized calcium signaling cellular organelles ^73–75^. However, the dependence of the function of primary cilia in nociceptor function on short and long range calcium signaling remains to be established.

Sensory transduction in nociceptors is modulated by diverse inflammatory mediators that sensitize nociceptors to produce mechanical hyperalgesia. Therefore, we also determined if primary cilia contribute to nociceptor sensitization by a clinically important pronociceptive inflammatory mediator, PGE_2_, whose synthesis is the target of the non-steroidal anti-inflammatory analgesic class of drugs (NSAIA/NSAID). In the present experiments treatment with an siRNA for *Ift88* markedly attenuated the mechanical hyperalgesia induced by PGE_2_. Again, how the primary cilium, located as a protrusion from the nociceptor cell body, which contains elements of signaling pathways implicated in nociceptor sensitization (e.g., adenylyl cyclase ^76–78^), contributes to sensitization of nociceptors, at their peripheral and central terminals, remains to be explored.

Since sensitized nociceptors play an important role in patients ^79–82^ and animals ^41,83–86^ with diverse forms of neuropathic pain, we also evaluated whether mechanical hyperalgesia associated with a preclinical model of CIPN, induced by a clinical neurotoxic cancer chemotherapy drug, paclitaxel, is also primary cilium dependent. Again, we observed that siRNA targeting *Ift88*, markedly attenuates the mechanical hyperalgesia associated with paclitaxel-induced CIPN.

To provide independent confirmation of the role of primary cilium in nociceptor function we evaluated the contribution of a second integral intra-flagellar protein, *Ift52*. As observed following i.t. treatment with siRNA for *Ift88*, siRNA for *Ift52* also markedly attenuated paclitaxel painful CIPN, supporting the suggestion that primary cilia contribute to nociceptor function. While details of the underlying mechanism by which the primary cilium signals to terminals of nociceptive sensory neurons remains to be elucidated, our results support a role of the primary cilium both in setting baseline nociceptive threshold and nociceptor sensitization associated with inflammation and peripheral neuropathy.

The primary cilium is a highly complex structure that contains many receptors, second messengers and ion channels^76,87,88^, some of which have been implicated in nociceptor function. One such primary cilium-dependent signaling pathway is Hh. Thus, it has been shown that selective inhibitors of Hh signaling (e.g., vismodegib and cyclopamine) attenuate mechanical hyperalgesia associated with inflammation^15,23^ and peripheral neuropathy ^19^, whereas activation of Hh signaling (e.g., with SAG, a smoothened agonist) has been found to produce hyperalgesia^19,23^. In the present study we demonstrate that the hyperalgesia induced by recombinant Shh is markedly attenuated in rats treated with siRNA for *Ift88*. Of note, however, in contrast to the effect of *Ift88* siRNA in control rats, which increased baseline mechanical nociceptive threshold in naïve control rats, inhibitors of Hh signaling in the absence of inflammation or peripheral neuropathy did not affect nociceptive threshold, supporting the presence of an Hh-independent primary cilium function capable of regulating baseline mechanical nociceptive threshold.

Another primary cilium-dependent cellular signaling pathway that has been found to contribute to nociception is the wingless-related integration site (Wnt)/μ-catenin signaling pathway ^89–95^. Ciliary proteins are not only necessary for the regulation of certain types of Wnt signaling but also include effectors downstream of Wnt ^96–99^. Of note, inhibitors of Wnt/μ-catenin signaling have been reported to attenuate vincristine- and paclitaxel-induced CIPN ^100–102^, and Wnt has been shown to be able to enhance neuronal excitability, producing peripheral sensitization ^103,104^. WNT agonists directly generate hypernociception ^105^; and, Wnt3a recruits Wnt-calcium signaling in sensory neurons to enhance pain sensitivity ^104,106^. Importantly, in contrast to Shh signaling, the fly homolog of μ-catenin, Armadillo, does regulate baseline sensitivity in nociceptors, in the absence of injury or inflammation ^91^. However, studies in species, in addition to *Drosophila*, and establishing whether this function of Wnt/μ-catenin signaling in nociceptor function is primary cilium dependent, and the mechanism by which it controls nociceptive threshold, remains to be established.

In summary, in the present study we have explored the role of the primary cilium and primary cilium-dependent Hh signaling, in nociceptors, in the regulation of baseline mechanical nociceptive threshold and inflammatory and neuropathic pain. We first established that primary cilia are present on multiple classes of primary afferent nociceptors in adult rats, *in vivo*, and in DRG neurons cultured from adult mice, *in vitro*. We next established that i.t. administration of siRNA for *Ift88*, an intra-flagellar protein essential for the integrity of the primary cilium, attenuates Ift88 in rat DRG and decreases both the number of DRG soma with primary cilia and the length of the remaining cilia. Treatment with *Ift88* siRNA produced a pain phenotype, including increased mechanical nociceptive threshold, and attenuation of inflammatory and neuropathic pain. To confirm the contribution of the primary cilium to the pain phenotype produced by siRNA targeting *Ift88*, we established a similar phenotype with siRNA targeting a second critically important intra-flagellar protein, *Ift52*. Thus, siRNA targeting multiple IFT genes coding proteins critical for integrity and function of primary cilia increased mechanical nociceptive threshold and attenuated inflammatory and neuropathic pain. Finally, we tested the hypothesis that primary cilium-dependent Hh signaling is involved in the role of the primary cilium in mechanical transduction in the nociceptor and its role in inflammatory and neuropathic pain. While primary cilium-dependent Hh signaling appears to play a role in nociceptor sensitization, in the absence of inflammatory or neuropathic pain it does not appear to contribute to mechanical nociceptive threshold. A major unresolved issue is how the primary cilium, which is located in the cell body of the primary afferent nociceptor, regulates the function of its remotely located central and peripheral terminals.

## Acknowledgements

All authors read and approved the final version of the manuscript. The authors thank Niloufar Mansooralavi for technical assistance and the UNE COBRE Histology and Imaging Core (NIGMS P20GM103643). This study was funded by National Institutes of Health (NIH) grants NS130249 (KLT), CA250017 (JDL) and AR075334 (JDL).

## Author contributions

L.S.-F., O.B., J.D.L., and K.L.T. conceived and designed the project. L.A.F., L.S.-F., O.B., D.A., I.J.M.B., E.E.J., and K.L.T. performed the experiments. L.S.-F., O.B., J.D.L., and K.L.T. wrote the manuscript.

## Conflict of interest statement

The authors declare no competing interests.

**Correspondence** and requests for materials should be addressed to Kerry L. Tucker

---

**Supplementary Figure 1.**
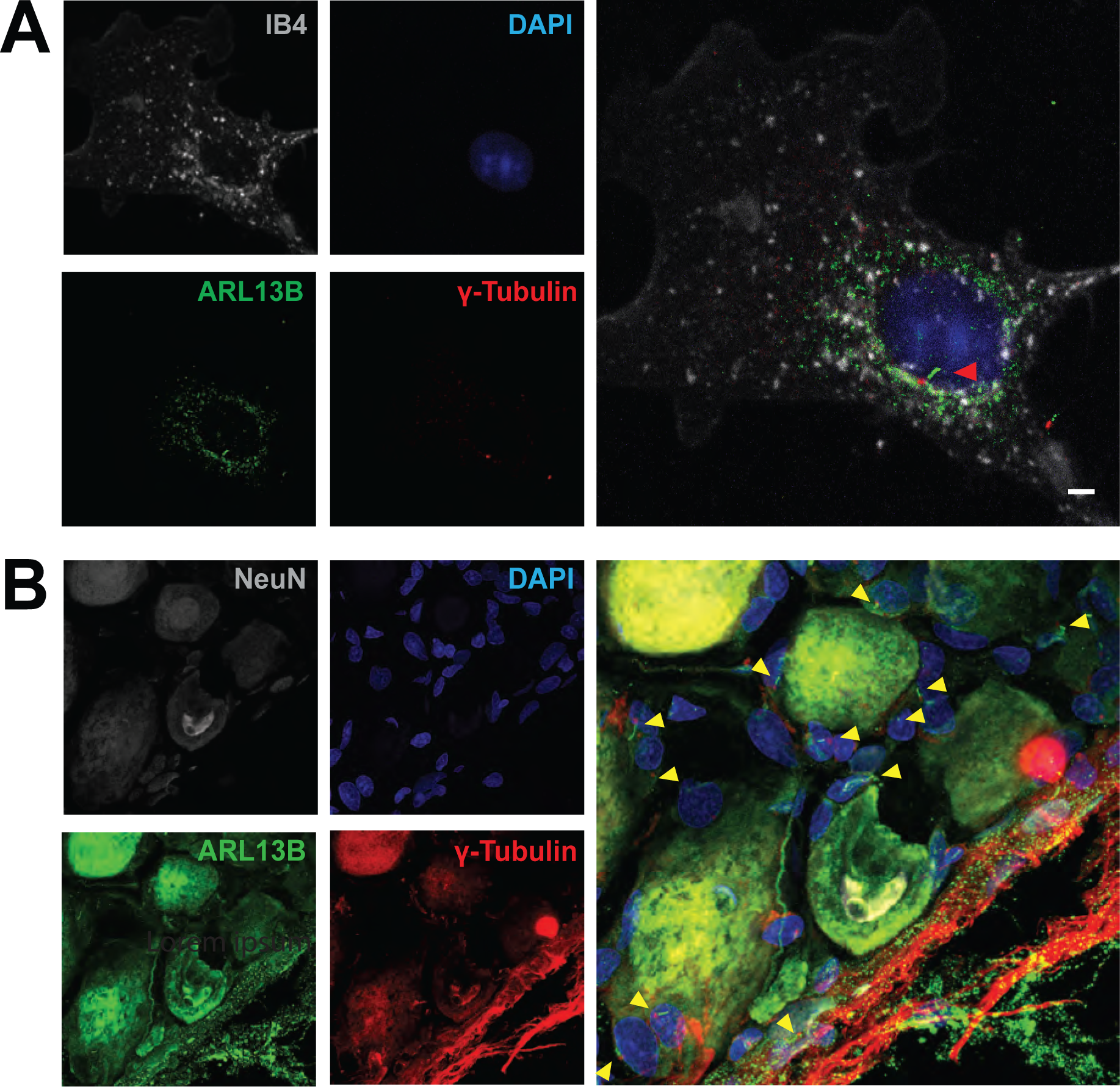
Mouse DRG neurons elaborate primary cilia *in vitro*, and satellite cells are also ciliated. (**A, B**) Immunohistofluorescence analysis of acutely dissociated adult mouse DRG culture (**A**) and of adult rat DRGs *in vivo* (**A, B**). Coverslips (**A**) and histological sections (**B**) were labeled with antibodies recognizing ARL13B (green), ɣ-tubulin (red), and Fox-3 (**B,** NeuN, greyscale). (**A**) IB4 staining is indicated in greyscale. (**A, B**) Cell nuclei marked by DAPI (blue). Red and yellow arrowheads indicate neuronal and non-neuronal primary cilia, respectively. Scale bar: 3 µm.

